# Exceptional biostability of paranemic crossover (PX) DNA, crossover-dependent nuclease resistance, and implications for DNA nanotechnology

**DOI:** 10.1101/801407

**Authors:** Arun Richard Chandrasekaran, Javier Vilcapoma, Paromita Dey, SiuWah Wong-Deyrup, Bijan K. Dey, Ken Halvorsen

## Abstract

Inherent nanometer-sized features and molecular recognition properties make DNA a useful material in constructing nanoscale objects, with alluring applications in biosensing and drug delivery. However, DNA can be easily degraded by nucleases present in biological fluids, posing a considerable roadblock to realizing the full potential of DNA nanotechnology for biomedical applications. Here we investigated the nuclease resistance and biostability of the multi-stranded motif called paranemic crossover (PX) DNA and discovered a remarkable and previously unreported resistance to nucleases. We show that PX DNA has more than an order of magnitude increased resistance to degradation by DNase I, serum, and urine compared to double stranded DNA. We further demonstrate that the degradation resistance decreases monotonically as DNA crossovers are removed from the structure, suggesting that frequent DNA crossovers disrupt either the binding or catalysis of nucleases or both. Further, we show using mouse and human cell lines that PX DNA does not affect cell proliferation or interfere with biological processes such as myogenesis. These results have important implications for building DNA nanostructures with enhanced biostability, either by adopting PX-based architectures or by carefully engineering crossovers. We contend that such crossover-dependent nuclease resistance could potentially be used to add “tunable biostability” to the many features of DNA nanotechnology.

## Introduction

Some daring purveyors of science fiction have imagined superior beings with more than 2 strands of DNA. In the 1997 movie *The Fifth Element*, for example, the heroine Leeloo is considered a perfect human-like being in part because of her 8-stranded helical DNA that is “tightly packed with infinite genetic knowledge”. While this is notably incorrect, there are instances both in biology and biotechnology where DNA structures can have more than two strands. Triplexes, for example can form a single helix using three strands,^1^ while guanine tetrads can form four-stranded DNA complexes.^2^

In DNA nanotechnology, synthetic DNA strands are designed and integrated together to form different motifs that serve as the building blocks for bottom-up construction.^3^ These (usually) multi-stranded structures typically contain double helical domains that are connected together by strand crossovers. A wide variety of structures have been made using bottom-up DNA construction, ranging from small objects and devices to larger, trigger responsive “cages”. While many DNA objects are seemingly made for their own sake (e.g. smiley face,^4^ bunny^5^), there are emerging applications of DNA nanotechnology such as drug delivery. To succeed as drug delivery vehicles, DNA objects must overcome a major challenge of surviving harsh “in vivo” environments such as blood.^6^ Strategies to improve the biostability of DNA structures include polymer coating, viral capsid encapsulation, modified nucleotides and crosslinking.^7^ One possibility that has been largely overlooked is that the inherent design of these DNA nanostructures can be altered to change biostability.

Construction of DNA nanostructures is based on robust starting units. One example is the double crossover (DX) motif that has two adjacent double helices connected by 2 crossover points.^8^ Design rules and construction parameters established on such DNA motifs are applied to other strategies and hierarchical assemblies (for example, the multi-crossover DNA origami).^4^ Another less common DNA motif is paranemic crossover (PX) DNA, a four-stranded DNA structure that consists of two adjacent and connected double helical DNA domains (**Figure 1a**).^9,10^ The motif is formed by creating crossovers between strands of the same polarity at every possible point between two side-by-side helices.^10^ Each duplex domain of PX DNA contains alternating major (wide) groove (denoted by W) or a minor (narrow) groove separation (denoted by N) flanking the central dyad axis of the structure, with one helical repeat containing a mixture of four half turns. Previous studies have reported PX DNA with different major/minor groove separations (W:N), with the most stable complexes containing 6, 7 or 8 nucleotides in the major groove and 5 nucleotides in the minor groove (PX 6:5, 7:5 and 8:5 respectively).^9,10^

**Figure 1.**
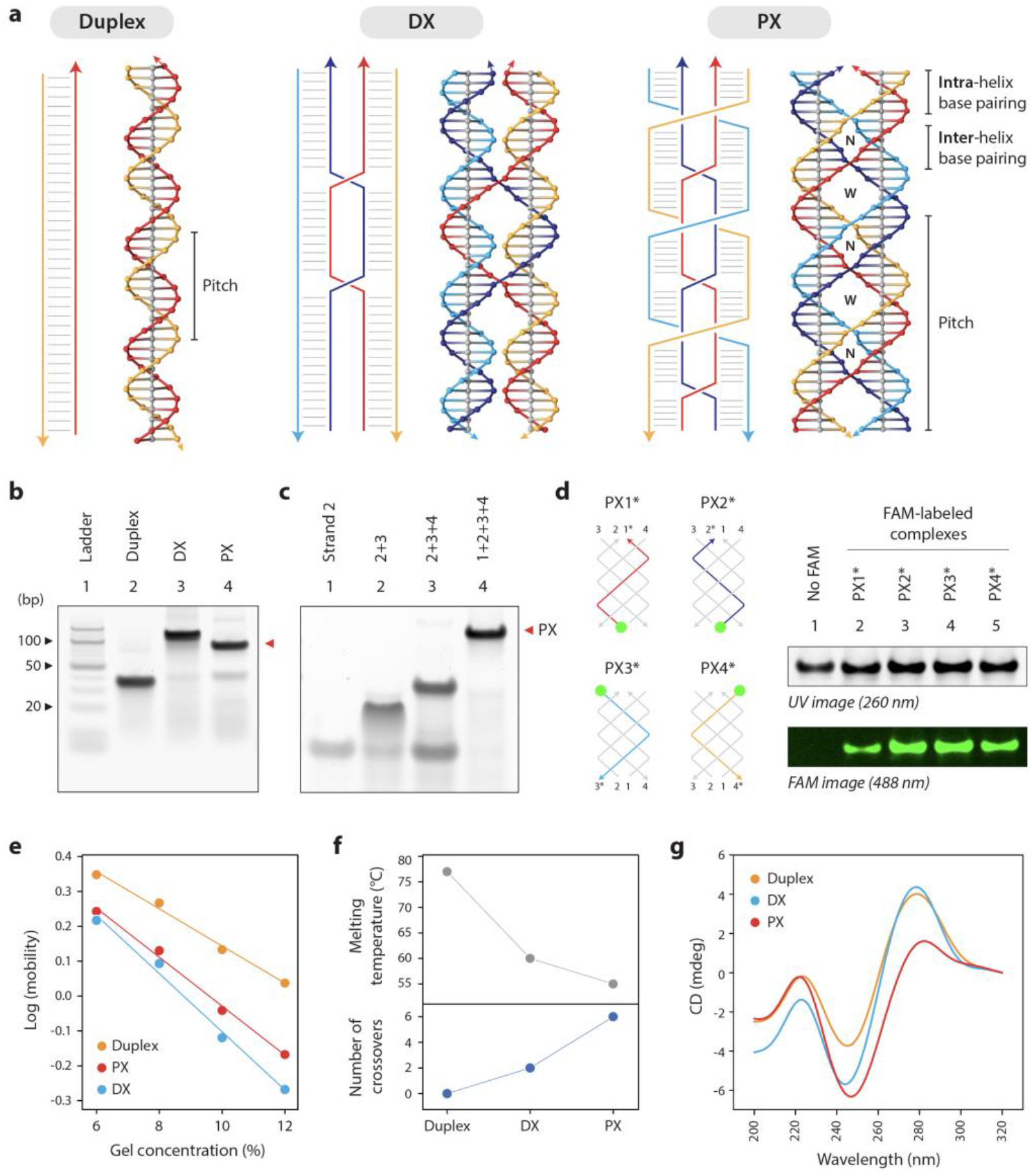
Design and characterization of paranemic crossover (PX) DNA. (a) Schematic and models of a B-DNA duplex, a double crossover (DX) motif and paranemic crossover (PX) motif. (b) Non-denaturing polyacrylamide gel showing formation of structures as predominant products in each lane, with PX migrating slightly faster than its DX counterpart. (c) Stepwise assembly of PX DNA with different numbers of strands in the complex. (d) Validating incorporation of all four strands in the PX. Strands 1, 2, 3 or 4 were labeled with a fluorescein (FAM, shown as green circles) and PX complexes containing one of these FAM labeled oligos were annealed. Gels imaged at the FAM-specific wavelength show that all four strands are present in the PX complex (lanes 2-5). Gel image under UV is shown for reference, with a control PX lane without any FAM labels (lane 1). (e) Ferguson plot showing gel mobility characteristics of the control and PX structures. (f) Melting temperatures determined from UV melting experiment show a decrease in thermal stability from duplex to DX to PX. (g) Circular dichroism spectra of the tested structures.

In DNA nanotechnology, PX DNA has been used to construct objects such as an octahedron^11^ and a triangle,^12^ as well as one- and two-dimensional arrays.^13,14^ PX DNA has also been a component of nanomechanical devices^15^ that are used in molecular assembly lines^16^ and DNA-based computation.^17^ In biology, PX DNA is studied for its involvement in double stranded DNA homology recognition due to its ability to relax supercoiled DNA.^18^ Recent studies also seek out proteins in the cell that can structure-specifically bind to PX DNA, so as to elucidate its biological function.^19,20^ These studies found that DNA polymerase I (Pol I) and T7 endonuclease I can bind to PX DNA, supporting the notion of its biological relevance. While PX DNA has been studied both in a nanotechnology and biological context, its biostability is largely unknown and its relative robustness compared to other motifs has not been analyzed in detail. In this work, we tested the nuclease resistance of PX DNA using DNase I and biostability of the motif in complex biological fluids such as human serum and urine.

## Results

In this study, we used a PX 6:5 molecule,^10,19^ where the helical repeat of each strand is 22 bp, making the pitch of the PX DNA motif roughly twice of that of B-DNA (with a net twist half of that of B-DNA) (**Figure 1a**, PX). We made two control structures; one DNA duplex with the sequence matching half of the PX motif (**Figure 1a**, **left**) and a double crossover (DX) motif with sequence similar to the PX but has only two crossover points (**Figure 1a**, **middle**). Figure 1 illustrates the crossover points in the strand diagrams for each structure and their respective molecular models (sequences are shown in **Figure S1**). We annealed the motifs in Tris-acetate-EDTA-Mg2+ buffer and checked their formation using non-denaturing polyacrylamide gel electrophoresis (PAGE) (**Figure 1b-c**). To further confirm the formation of the four-stranded PX, we labeled each of the four strands with fluorescein (FAM). We then annealed PX complexes where only one of the four component strands were labeled with FAM (**Figure 1d and Figure S2**). The presence of a fluorescent band in each of these four lanes on a non-denaturing PAGE indicated that all four strands are present in the complex (lanes 2-5) compared to a non-labeled complex (lane 1). Next, we analyzed the electrophoretic mobility of the different motifs as a function of gel concentration using a Ferguson plot (**Figure 1e** and **Figure S3**), where the slope can be used to estimate the retardation (frictional) coefficient of the structures. The plot shows that the slope of the PX molecule is comparable to that of the DX, and distinct from a regular double stranded DNA, a trend that is consistent with previous results.^10^ We then analyzed the thermal melting profiles of the structures and obtained the melting temperatures of the duplex (77 °C), DX (60 °C) and PX (55 °C) (**Figure 1f** and **Figure S4**). It can be seen that the thermal stability decreases with higher number of crossovers in the structures. Circular dichroism profiles of the duplex, DX and PX were also consistent with previous reports available for these structures (**Figure 1g**).

To investigate biostability, we first tested the nuclease resistance of PX DNA using DNase I, an endonuclease that nonspecifically cleaves double stranded DNA.^21^ To perform our DNase I assay at the optimal and physiologically relevant temperature of 37 °C, we first confirmed that the structures were intact at this elevated temperature. We incubated the duplex, DX and PX DNA at 37 °C for 24 hours and showed that physiological temperature does not noticeably affect the structures in this time (**Figure S5**). We annealed the duplex and DX controls and the PX DNA, and incubated them with various amounts of DNase I enzyme for different times at 37 °C. We ran the DNase I treated samples on non-denaturing PAGE and quantified the reduction in the band corresponding to the structures to obtain the intact fraction for each structure (**Figure 2a** and **Figure S6**). With 0.1 units of DNase I, the duplex and DX structures were almost fully degraded within a few minutes (85% duplex and 70% DX degraded in 1 min), while PX DNA remained largely intact even after 1 hour. When we increased the DNase I to 0.25 units (and up to 1 unit), both the duplex and DX motif were degraded completely in under 2 mins (**Figure S6**). To observe complete degradation of PX DNA by 1 hour, we had to increase the DNase I amount to 1 unit (**Figure 2b**). Using this DNase I concentration, we checked the cleaved products from PX at different incubation times with the enzyme. We annealed the PX with one of the component strands labeled with FAM, and ran the DNase I treated samples on a denaturing gel (**Figure S7**). We imaged the gel at the wavelength for FAM and quantified the FAM-labeled single strand (**Figure 2c**). By the 2-minute time point in the DNase I reaction, 60% of the products were shorter than the single strand (38 nt) and by 16 mins, there was no single stranded product remaining and all of the complex was digested into polynucleotides. On comparing the structures at different DNase I amounts, PX DNA was at least 10-fold more resistant to nuclease degradation than duplex or DX (**Figure 2d**).

**Figure 2.**
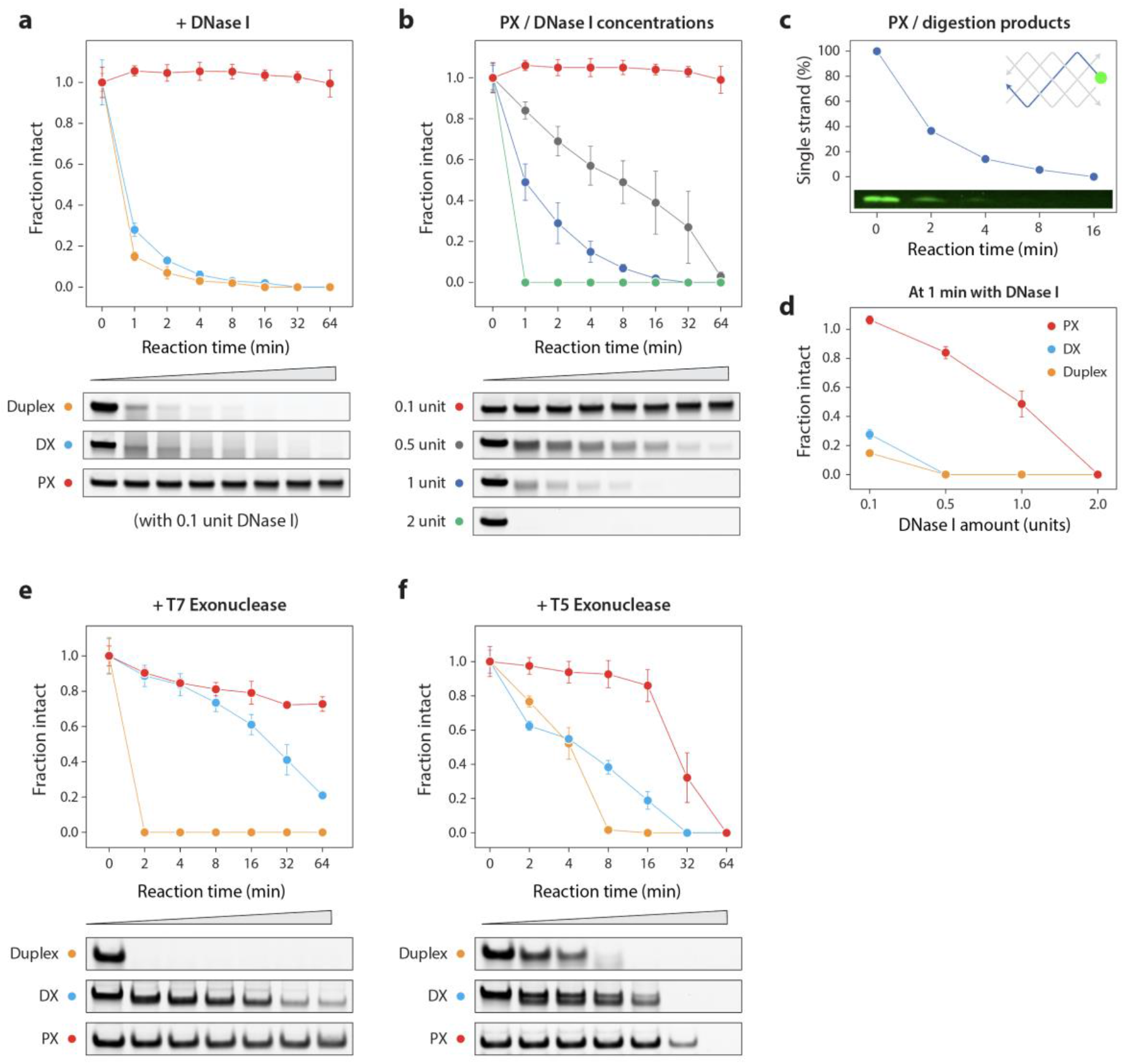
Nuclease resistance of duplex, DX and PX DNA. (a) Degradation plot of the structures when treated with 0.1 unit of DNase I enzyme. Non-denaturing gels containing structures at different time points with DNase I are shown below the plot. (b) Quantitative plot and non-denaturing gels of PX DNA treated with different amounts of DNase I. (c) Digestion of PX monitored by fluorescently labeled component strand. (d) Comparative degradation of duplex, DX and PX DNA in different DNase I amounts at the 1 min time point. Degradation plots and non-denaturing gels of structures treated with (e) T7 exonuclease and (f) T5 exonuclease.

Next, we tested whether this nuclease resistance of PX is true beyond DNase I. We incubated the annealed PX complex with two other exonucleases and tested digestion at different time points. Similar to the trends with DNase I, PX was the most stable in the reactions with T7 exonuclease (**Figure 2e**) and T5 exonuclease (**Figure 2f**) compared to control duplex and DX structures (full gels in **Figure S8**).

Considering the unusual nuclease resistance of PX DNA, we hypothesized that the nuclease may be obstructed by the frequent crossovers. To test this hypothesis, we designed and constructed variations of PX DNA that have sequentially fewer crossover points. Such structural variations were previously termed juxtaposed crossover (JX) DNA motifs. The JX motifs were further denoted with numbers signifying the missing crossovers compared to PX structure (e.g. JX2 denotes two missing crossovers) (**Figure 3a** and **Figure S9**).

**Figure 3.**
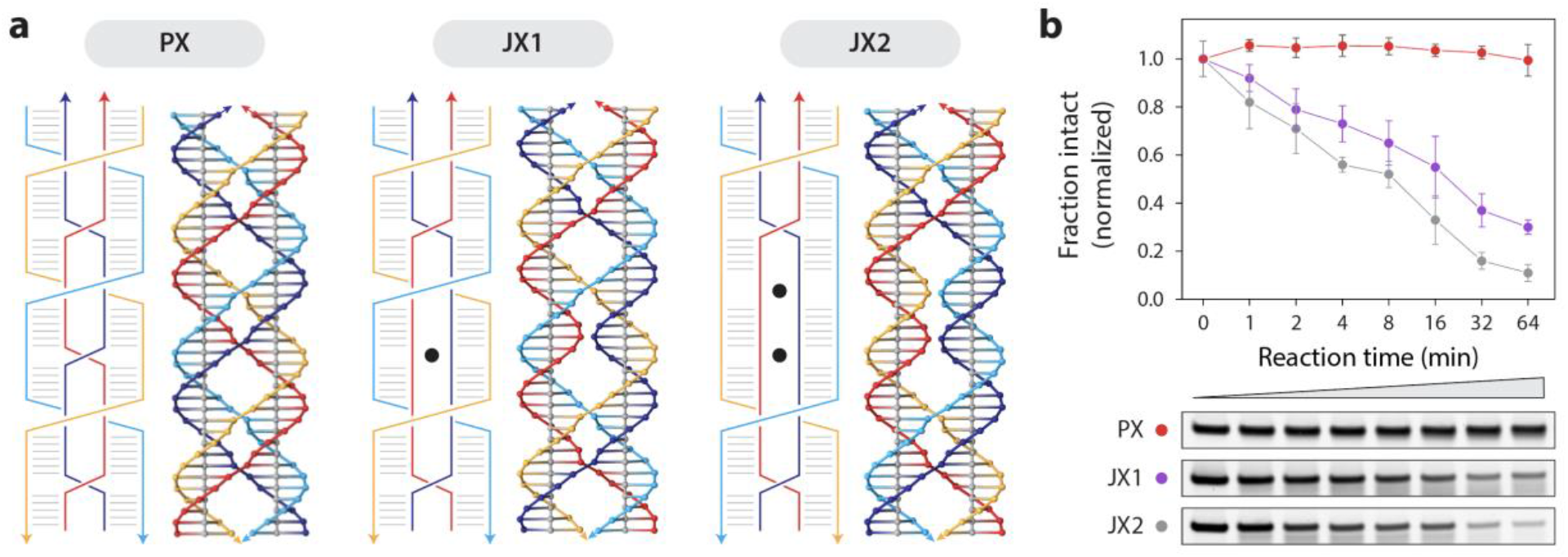
Characterization and nuclease resistance of PX and JX structures. (a) Schematic and models of the PX motif and its variations JX1 and JX2 that lack 1 and 2 crossovers respectively (shown as black dots in the structural diagram). (b) Degradation plot and non-denaturing gels of the PX, JX1 and JX2 structures when treated with 0.1 unit of DNase I enzyme.

Based on the PX, we designed strands to make JX motifs JX1, JX2 and JX3. Following similar protocols for formation of PX, we successfully formed all three structures as seen on non-denaturing PAGE (**Figure S10**). To our knowledge, the JX1 and JX2 structures have been experimentally demonstrated in the lab,^15,22^ while the other versions have been simulated by molecular dynamics and predicted to be stable to varying degrees.^23,24^ However, when we checked their stability at 37 °C we found that the JX3 structure was not stable (**Figure S11)** and so we omitted this structure from further experiments.

We incubated the JX1 and JX2 structures with 0.1 unit DNase I for different time periods and tested degradation of the structures on non-denaturing PAGE (**Figure 3b and Figure S12**). Quantified results of the DNase I assay shown in **Figure 3b** demonstrates that PX DNA is the most stable, and the order of DNase I resistance is PX > JX1 > JX2. These results indicate that structures with more crossovers (PX) are more resistant to DNase I activity, supporting our original hypothesis.

To further investigate the biostability of these structures, we tested complex biological fluids such as fetal bovine serum (FBS), human serum and human urine. We incubated the duplex, DX, PX, JX1 and JX2 motifs in 10% FBS for various time points up to 24 hours at 37 °C, and analyzed the treated samples on non-denaturing PAGE (**Figure 4a** and **Figure S13**). As seen with the DNase I assay, quantified results showed that PX DNA was again the most stable, with no discernable degradation in 10% FBS even after 24 hours, while the duplex and DX were completely degraded at the 24-hour time point (plot in **Figure 4a**). The JX1 and JX2 structures were partially degraded at the 24-hour time point (25% and 74% degraded, respectively). We also tested the duplex, DX and PX DNA in different percentages of FBS for 1.5 hours and PX was more stable in all cases (**Figure S14**). To ascertain the biological relevance in human context, we then tested how the biostability of these structures compare in human serum and urine. We incubated the structures in 10% serum and 10% urine for 24 hours at 37 °C (**Figure 4b** and **Figure S15**). In both human serum and urine, the PX remained intact after 24 hours while the other structures showed various increased levels of degradation (**Figure 4b**).

**Figure 4.**
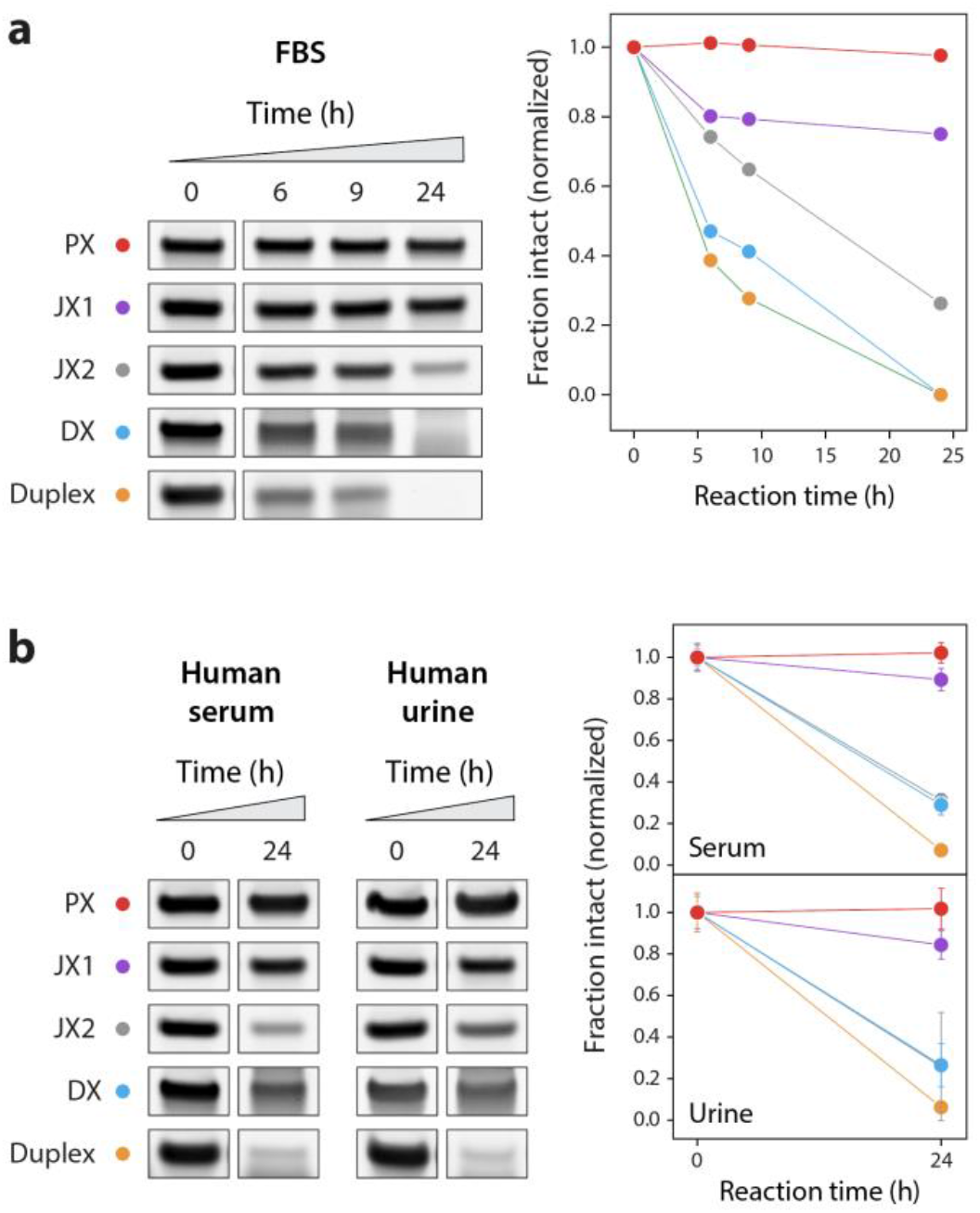
Stability in complex biological fluids. (a) Non-denaturing gels and degradation plot of control structures (duplex and DX) and PX, JX1, and JX2 in 10% fetal bovine serum (FBS). (b) Stability of the structures when incubated in 10% human serum and human urine for 24 hours. Non-denaturing gels show the bands corresponding to initial structures and at 24 hours. Quantitative plot for all structures in serum and urine is shown on the right.

From the results of nuclease resistance and biofluids experiments, we did a comparative analysis of all the tested structures to find out the structural and biostability trend as a function of the number of crossovers in the structure (**Figure 5**). We quantified the intact fraction of each structure in 0.1 unit DNase I (grey), 10% FBS (blue), 10% human serum (red) and 10% human urine (yellow). The overall stability of the structures in all these conditions was PX > JX1 > JX2 > DX > duplex (with 6, 5, 4, 2 and 0 crossover points respectively), reflecting the effect of crossovers on nuclease resistance and biostability.

**Figure 5.**
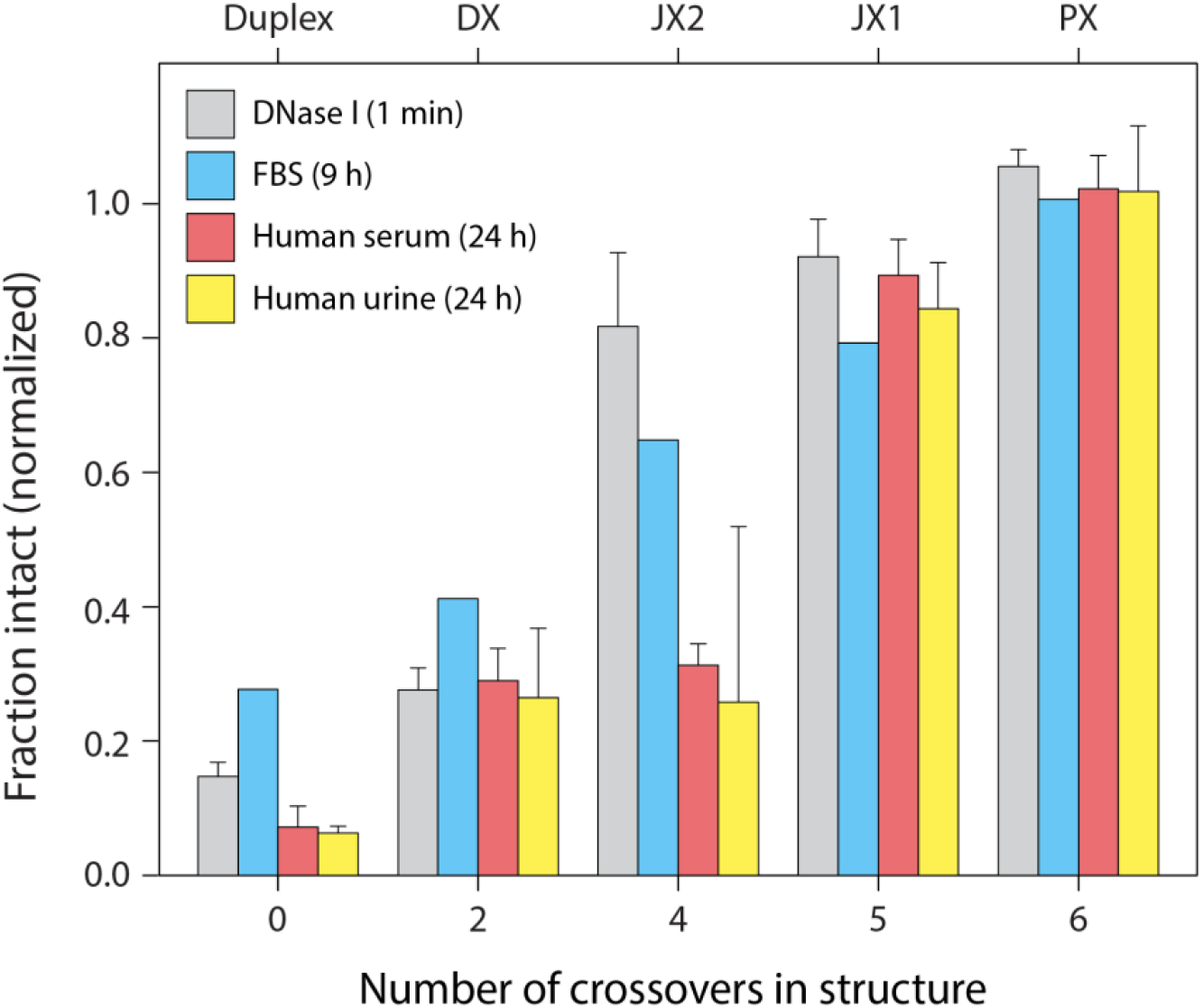
Crossover number vs biostability. A comparative analysis of the tested structures (duplex, DX, PX, JX1 and JX2) in DNase I enzyme (at 1 min reaction time), 10% FBS (at 9 hours), 10% human serum (at 24 hours) and 10% human urine (at 24 hours). Results show an upward trend in nuclease resistance and biostability with increasing number of crossovers in the structure (PX > JX1 > JX2 > DX > duplex).

The exceptional biostability of PX DNA indicates potential use of this motif to construct drug delivery carriers. To validate the potential use of PX-based structures in drug delivery, we tested cell viability using MTT assay in mouse (C2C12) and human (HeLa) cell lines. After testing different concentrations of PX incubated with these cells (**Figure S16**), we analyzed cell viability after 24, 48 and 72 hours of incubation with PX (**Figure 6a-b**). We further showed that PX does not interfere with cellular processes by incubating differentiating skeletal muscle cells with PX. We cultured cells in differentiation medium and observed cell differentiation from growth medium (undifferentiated) and after 2 and 4 days in differentiation medium (differentiated cells, DM2 and DM4) (**Figure 6c**). We confirmed differentiation in these cells qRT-PCR analysis of early and late myogenic differentiation markers myogenin (Myog, **Figure 6d**) and Myosin Heavy Chain (MHC, **Figure 6e**) respectively. Results showed that these biomarkers were increased with the same fold-change in PX-incubated cells as compared to the control cells without PX.

**Figure 6.**
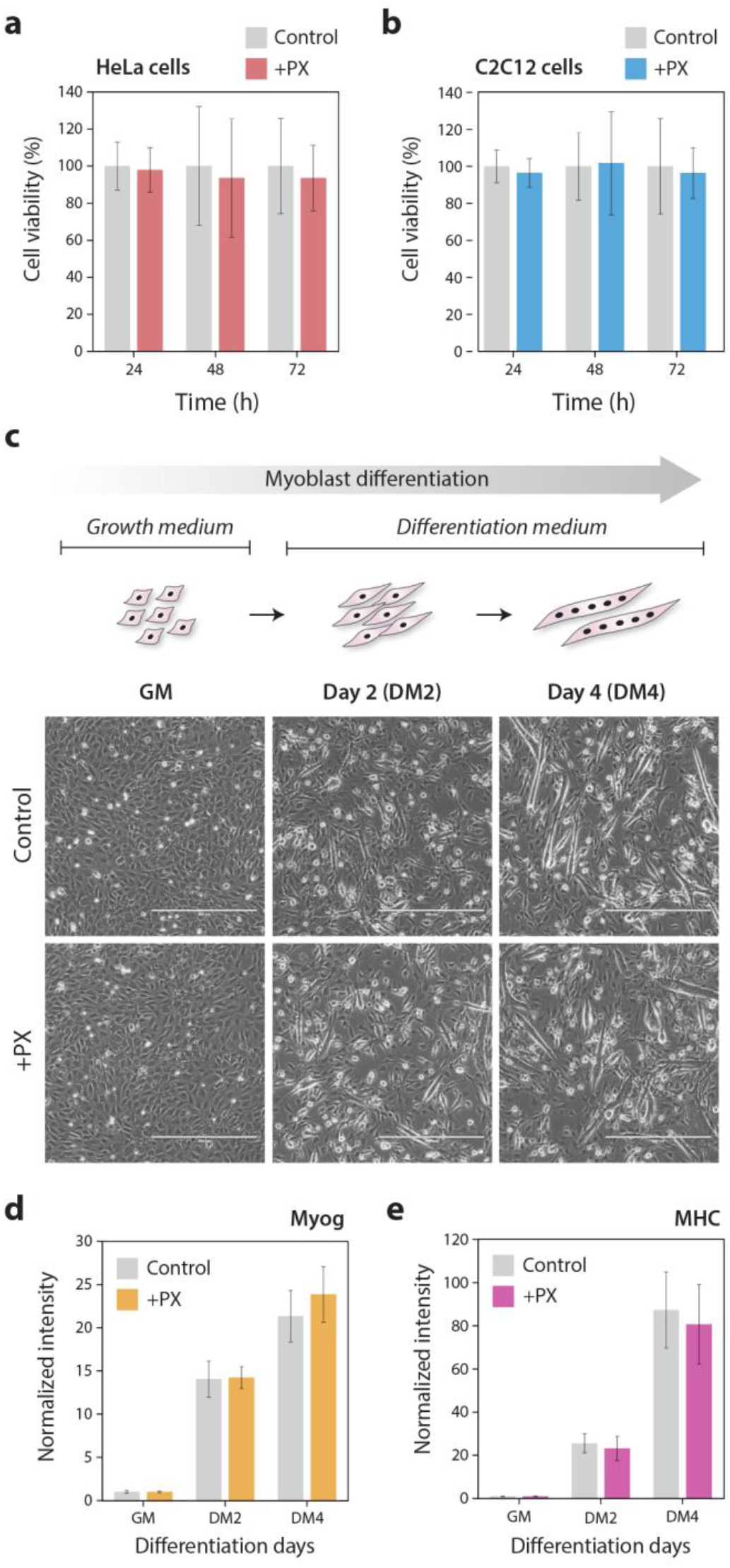
PX DNA does not affect cell viability or influence differentiation of skeletal muscle cells. (a) and (b) show cell viability from MTT assay when HeLa cells and C2C12 cells were incubated with 100 nM PX DNA for 24-72 hours. (c) Addition of PX DNA does not affect differentiation of myoblast cells. Images show undifferentiated cells (GM) and myocytes in early (DM2) and late (DM4) differentiation stages. Scale bars are 400 μm. (d) and (e) are qRT-PCR results of myogenic differentiation markers Myog and MHC both of which are upregulated in DM1 and DM4 cells in both the control set and cells incubated with PX DNA.

## Discussion

Our results clearly demonstrate for the first time that PX DNA has a dramatically enhanced biostability compared to normal double stranded DNA. Quantitatively, this effect is at least an order of magnitude difference for all the tests we performed. This remarkable difference could have a number of plausible (and not mutually exclusive) explanations that may merit further study. Enzymatic activity on DNA is known to depend on the helical twist of DNA molecules,^25^ with DNase I in particular dependent on groove width and flexibility of the duplex.^26^ Thus, the enhanced resistance of PX DNA to DNase I could be in part due to the difference in helical parameters compared to that of regular DNA duplexes. DNase I has also been previously shown to require a substrate of typically 6-8 base pairs,^26,27^ and digestion of branched junctions has shown that several nucleotides near the branch point are protected from DNase I cleavage.^28^ In PX DNA, crossovers occur every half turn and thus the available double helical region between consecutive crossovers is only 5 or 6 bps (alternating half turns). These small regions may make DNase I binding difficult or impossible and slow the degradation. Our results on the JX motifs are consistent with these possible explanations, as removal of each crossover will both extend potential binding regions and relax the DNA in that region.

Previous studies have shown that DNA nanostructures exhibit enhanced resistance to nucleases compared to oligonucleotides or plasmid DNA.^29^ Studies have also shown that DNA origami structures incubated in tissue culture medium containing 10% FBS showed degradation within two hours.^30^ There are some strategies available to overcome these effects and increase the lifetime of DNA nanostructures in biofluids. These include heat treatment of FBS,^30^ addition of actin protein that inhibits nuclease activity,^30^ or chemical modification of component DNA strands.^31^ However, these strategies are not without disadvantages. Heat treatment and addition of external proteins can affect the physiological environment and are probably not feasible *in vivo*, while chemically modified nucleic acids can sometimes be toxic or induce unwanted immune responses.

The exceptional nuclease resistance of PX DNA suggests that it may be possible to design DNA nanostructures that could be inherently more biostable. While it is difficult to make direct quantitative comparison against previous studies, it is worth noting that our PX DNA showed no signs of degradation at 24 hours in 10% FBS, while DNA origami objects were largely destroyed under similar conditions.^30^ Noting that DNA nanostructures tend to have enhanced nuclease resistance compared to their individual structural components, we predict that larger DNA objects constructed from PX motifs will dramatically outlive their DNA origami counterparts. While a few DNA nanostructures have utilized PX DNA before, the relative biostability of PX nanostructures and non-PX versions have not been investigated. For example, the recently developed single-stranded origami^32^ approach utilizes the PX structure; it would be interesting to test the biostability of these structures compared to “regular” origami structures to see if our predictions hold true. Moreover, our study shows that incubation with PX does not interfere with cellular processes such as differentiation, indicating potential use of PX-based nanostructures in *in vivo* applications.

Beyond using PX DNA to increase robustness for biological applications of DNA nanostructures, our discovery of crossover-dependent biostability suggests that biostability can be designed into DNA nanostructures. For example, it could be possible to add protection to structures by strategic placement of crossovers at areas that are especially exposed to nucleases. Such a strategy, if successful, could lead to DNA nanostructures with a “tunable” biostability dictated by bottom up design principles. In drug delivery applications, this type of tailored biostability would facilitate timed degradation for fast or slow release of the encapsulated cargo. Further studies in DNA nanostructures containing different number or arrangement of crossovers are certainly needed to see if such speculation is grounded in dream or reality. The sci-fi writers of *The Fifth Element* were not always (or ever) grounded in reality, but they were inadvertently on to something with enhancement by multiple DNA strands. At least for the 4 stranded PX DNA structure, perhaps enhanced biostability is “the fifth element”.

## Acknowledgements

Research reported in this publication was supported by the NIH through NIGMS under award R35GM124720 to K.H. and by the American Heart Association under award 17SDG33670339 to B.K.D. We thank Dr. Jibin Abraham Punnoose for discussions on the project, Johnsi Mathivanan for assistance with CD experiments, and Andrew Hayden for the pop-culture reference to multi-stranded DNA in *The Fifth Element*.

## Author contributions

ARC conceived and designed the project. ARC and KH designed experiments. ARC, JV and SWD performed characterization experiments. ARC and JV performed in vitro nuclease resistance and biological fluid experiments. PD and BKD carried out cell culture, myoblast differentiation and MTT assays. ARC wrote initial draft of the paper. ARC and KH edited later drafts of the paper.

## References

1. Chandrasekaran, A. R. & Rusling, D. A. Triplex-forming oligonucleotides: a third strand for DNA nanotechnology. Nucleic Acids Res. 46, 1021–1037 (2018).

2. Lipps, H. J. & Rhodes, D. G-quadruplex structures: in vivo evidence and function. Trends Cell Biol. 19, 414–422 (2009).

3. Seeman, N. C. & Sleiman, H. F. DNA nanotechnology. Nat. Rev. Mater. 3, 17068 (2018).

4. Rothemund, P. W. K. Folding DNA to create nanoscale shapes and patterns. Nature 440, 297–302 (2006).

5. Benson, E. et al. DNA rendering of polyhedral meshes at the nanoscale. Nature 523, 441–444 (2015).

6. Madhanagopal, B. R., Zhang, S., Demirel, E., Wady, H. & Chandrasekaran, A. R. DNA Nanocarriers: Programmed to Deliver. Trends Biochem. Sci. 43, 997–1013 (2018).

7. Kizer, M. E., Linhardt, R. J., Chandrasekaran, A. R. & Wang, X. A Molecular Hero Suit for In Vitro and In Vivo DNA Nanostructures. Small 15, 1805386 (2019).

8. Fu, T. J. & Seeman, N. C. DNA double-crossover molecules. Biochemistry 32, 3211–3220 (1993).

9. Wang, X. et al. Paranemic Crossover DNA: There and Back Again. Chem. Rev. 119, 6273–6289 (2019).

10. Shen, Z., Yan, H., Wang, T. & Seeman, N. C. Paranemic Crossover DNA: A Generalized Holliday Structure with Applications in Nanotechnology. J. Am. Chem. Soc. 126, 1666–1674 (2004).

11. Shih, W. M., Quispe, J. D. & Joyce, G. F. A 1.7-kilobase single-stranded DNA that folds into a nanoscale octahedron. Nature 427, 618 (2004).

12. Liu, W., Wang, X., Wang, T., Sha, R. & Seeman, N. C. PX DNA Triangle Oligomerized Using a Novel Three-Domain Motif. Nano Lett. 8, 317–322 (2008).

13. Ohayon, Y. P. et al. Covalent Linkage of One-Dimensional DNA Arrays Bonded by Paranemic Cohesion. ACS Nano 9, 10304–10312 (2015).

14. Shen, W. et al. The study of the paranemic crossover (PX) motif in the context of self-assembly of DNA 2D crystals. Org. Biomol. Chem. 14, 7187–7190 (2016).

15. Yan, H., Zhang, X., Shen, Z. & Seeman, N. C. A robust DNA mechanical device controlled by hybridization topology. Nature 415, 62 (2002).

16. Gu, H., Chao, J., Xiao, S.-J. & Seeman, N. C. A proximity-based programmable DNA nanoscale assembly line. Nature 465, 202–205 (2010).

17. Chakraborty, B., Jonoska, N. & Seeman, N. C. A programmable transducer self-assembled from DNA. Chem. Sci. 3, 168–176 (2011).

18. Wang, X., Zhang, X., Mao, C. & Seeman, N. C. Double-stranded DNA homology produces a physical signature. Proc. Natl. Acad. Sci. 107, 12547–12552 (2010).

19. Gao, X. et al. The PX Motif of DNA Binds Specifically to Escherichia coli DNA Polymerase I. Biochemistry 58, 575–581 (2019).

20. Kizer, M. et al. Complex between a Multicrossover DNA Nanostructure, PX-DNA, and T7 Endonuclease I. Biochemistry 58, 1332–1342 (2019).

21. Laskowski, M. 12 Deoxyribonuclease I. in The Enzymes (ed. Boyer, P. D.) 4, 289–311 (Academic Press, 1971).

22. Spink, C. H., Ding, L., Yang, Q., Sheardy, R. D. & Seeman, N. C. Thermodynamics of Forming a Parallel DNA Crossover. Biophys. J. 97, 528–538 (2009).

23. Maiti, P. K., Pascal, T. A., Vaidehi, N. & Goddard, W. A. The stability of Seeman JX DNA topoisomers of paranemic crossover (PX) molecules as a function of crossover number. Nucleic Acids Res. 32, 6047–6056 (2004).

24. Santosh, M. & Maiti, P. K. Structural Rigidity of Paranemic Crossover and Juxtapose DNA Nanostructures. Biophys. J. 101, 1393–1402 (2011).

25. Lomonossoff, G. P., Butler, P. J. G. & Klug, A. Sequence-dependent variation in the conformation of DNA. J. Mol. Biol. 149, 745–760 (1981).

26. Weston, S. A., Lahm, A. & Suck, D. X-ray structure of the DNase I-d(GGTATACC)2 complex at 2·3Å resolution. J. Mol. Biol. 226, 1237–1256 (1992).

27. Suck, D. DNA recognition by DNase I. J. Mol. Recognit. 7, 65–70 (1994).

28. Lu, M., Guo, Q., Seeman, N. C. & Kallenbach, N. R. DNase I cleavage of branched DNA molecules. J. Biol. Chem. 264, 20851–20854 (1989).

29. Keum, J.-W. & Bermudez, H. Enhanced resistance of DNAnanostructures to enzymatic digestion. Chem. Commun. 7036–7038 (2009). doi:10.1039/B917661F

30. Hahn, J., Wickham, S. F. J., Shih, W. M. & Perrault, S. D. Addressing the Instability of DNA Nanostructures in Tissue Culture. ACS Nano 8, 8765–8775 (2014).

31. Conway, J. W., McLaughlin, C. K., Castor, K. J. & Sleiman, H. DNA nanostructure serum stability: greater than the sum of its parts. Chem. Commun. 49, 1172–1174 (2013).

32. Han, D. et al. Single-stranded DNA and RNA origami. Science 358, eaao2648 (2017).

