## Supplementary Information for "Exceptional biostability of paranemic crossover (PX) DNA, crossover-dependent nuclease resistance, and implications for DNA nanotechnology"

**Materials and methods**

### Oligonucleotide sequences

All oligonucleotides were purchased from IDT, Inc. Full sequences (5'-3') are listed below. Strand routing for duplex, DX and PX structures are shown in Figure S1, and $JX_1$, $JX_2$ and $JX_3$ in Figure S9.

DC1A: 5’-GTGGTGTCGCAACAGATATGTGTAGGCACCGAATCACT-3’

DC1B: 5’-AGTGATTCGGTGCCTACACATATCTGTTGCGACACCAC-3’

DX1: 5’-AGTGATTCGGTGCCTACACATATCTGTTGCGACACCAC-3’

DX2: 5’-GTGGTGTCGCAACAGACACAATACTTCACCGAATCACT-3’

DX3: 5’-ACTAGATCATCAATGCTATGTGTAGGGTTAGACCTGAG-3’

DX4: 5’-CTCAGGTCTAACAAGTATTGTGGCATTGATGATCTAGT-3’

PX1: 5’-GTGGTATCATCAATGCTATGTGTAGGCTTAGACCTGAG-3’

PX2: 5’-ACTAGGTCGCAACAGACACAATACTTGACCGAATCACT-3’

PX3: 5’-AGTGAGTCTAACAAGTCACATATCTGTGATGATCTAGT-3’

PX4: 5’-CTCAGTTCGGTGCCTAATTGTGGCATTTGCGACACCAC-3’

JX-A: 5’-GTGGTATCATCAATGCCACAATACTTGACCGAATCACT-3’

JX-B: 5’-ACTAGGTCGCAACAGATATGTGTAGGCTTAGACCTGAG-3’

JX-C: 5’-AGTGAGTCTAACAAGTATTGTGGCATTTGCGACACCAC-3’

JX-D: 5’-CTCAGTTCGGTGCCTACACATATCTGTGATGATCTAGT-3’

JX-E: 5’-AGTGAGTCTAACAAGTATTGTGGCATTGATGATCTAGT-3’

JX-F: 5’-CTCAGTTCGGTGCCTACACATATCTGTTGCGACACCAC-3’

Fluorophore-labeled strands:

PX1: 5’-FAM-GTGGTATCATCAATGCTATGTGTAGGCTTAGACCTGAG-3’

PX2: 5’-FAM-ACTAGGTCGCAACAGACACAATACTTGACCGAATCACT-3’

PX3: 5’-FAM-AGTGAGTCTAACAAGTCACATATCTGTGATGATCTAGT-3’

PX4: 5’-FAM-CTCAGTTCGGTGCCTAATTGTGGCATTTGCGACACCAC-3’

### Formation of DNA complexes

To form the different DNA complexes (duplex, DX, PX and JXn), we mixed the specific component strands (Figure S1) in equimolar ratios to a final concentration of 1 μM in Tris-Acetic-EDTA-$Mg^{2+}$ (TAE/$Mg^{2+}$) buffer containing 40 mM Tris base (pH 8.0), 20 mM acetic acid, 2 mM EDTA, and 12.5 mM magnesium acetate. The DNA solution was slowly cooled from 95°C to 20 °C by placing the tubes in 2L of hot water in a styrofoam box for 48 hours to facilitate hybridization.

**DNase I assay**

Annealed DNA complexes (at 1 μM) were first mixed with DNase I reaction buffer (final of 1X). Dilutions of DNase I enzyme (NEB) to different units was made in nuclease-free water. For the DNase I assay, 1 μL of the enzyme was added to 10 μL of the sample containing DNase I reaction buffer. Samples were incubated at 37 °C for different time intervals. Incubated samples were mixed with gel loading dye containing bromophenol blue and 1X TAE/$Mg^{2+}$ and run on non-denaturing PAGE to analyze degradation over time.

**Stability test in biofluids**

Pooled normal human serum and human urine were purchased from Innovative Research Inc. Fetal bovine serum (FBS), human serum or human urine was added to annealed DNA complexes to be at a final concentration of 10%. Typically, we added 1μL of the biofluid to 10 μL of the annealed samples and incubated them at 37 °C for different time intervals. Incubated samples were mixed with gel loading dye containing bromophenol blue and 1X TAE/$Mg^{2+}$ and run on non-denaturing PAGE to analyze degradation over time.

**Native polyacrylamide gel electrophoresis (PAGE)**

Non-denaturing gels containing 4% polyacrylamide (29:1 acrylamide/bisacrylamide) were run at 4°C (100 V, constant voltage) in 1X TAE/$Mg^{2+}$ running buffer. After electrophoresis, the gels were stained in 1X TAE/$Mg^{2+}$ buffer containing 0.5X GelRed (Sigma). Imaging was done on a Bio-Rad Gel Doc XR+ imager using the default settings for GelRed with UV illumination. Quantification was done using the highest exposure image that did not contain saturated pixels in the band of interest. Gel images were exported as 12-bit images and quantified using ImageJ.

**UV melting experiments**

Duplex, DX and PX complexes were annealed at 1 μM concentration and used for UV melting experiments. Experiments were performed in a Cary 300 UV-Visible Spectrophotometer equipped with a temperature controller. Melting curves were acquired at 260 nm by heating and cooling from 5 °C to 85 °C at a rate of 0.2 °C/min.

**Circular dichroism experiments**

CD spectra was collected on samples annealed in 1X TAE/$Mg^{2+}$ at 5 μM concentration. Experiments were performed on a Jasco-815 CD spectrometer at room temperature in a quartz cell with a 10-mm path length. CD spectra were collected from 380 to 200 nm and with a scanning speed of 100 nm/min. The bandwidth was 1.0 nm, and the digital integration time was 1.0 s.

**Cell culture and differentiation assay**

Mouse myoblast cell line (C2C12) and human cell line (HeLa) were acquired from the American Type Culture Collection (ATCC). Cells were maintained at subconfluent densities in GM at 37 °C in a tissue culture incubator with a constant supply of 5% $CO_2$. GM consists of Dulbecco's modified Eagle medium (DMEM; Gibco) supplemented with 10% FBS and 1× antibiotic- antimycotic (Life Technologies). For myogenic differentiation assay, the myoblast cells were grown to about 70% confluency, washed with phosphate-buffered saline (PBS), and cultured with DM. DM consists of DMEM containing 2% heat-inactivated horse serum (HyClone) and 1× antibiotic-antimycotic. Cells were harvested while growing in GM and after 48 and 96 hours (DM2 and DM4, respectively) in DM. For the control set, cells were incubated with 1X TAE and for the test set, cells were incubated with 100 nM PX DNA dissolved in 1X TAE.

**MTT assay**

MTT assay was carried out using Vybrant® MTT Cell Proliferation Assay Kit (Invitrogen) following the manufacturer's instruction. Briefly, the medium was removed and replace it with 100 µL of fresh culture medium without phenol red. 10 µL of the 12 mM MTT stock solution was then added to each well and incubated at 37°C for 4 hours. After labeling the cells with MTT, all the medium and reagent mixtures were removed but 25 µL of medium and reagent mixtures were left in each well. Then, 50 µL of DMSO was added to each well, mixed thoroughly and incubated at 37°C for 10 minutes. The samples were mixed well once again and absorbance was read at 540 nm using a plate reader. For the control set, cells were incubated with 1X TAE and for the test set, cells were incubated at the indicated time with various concentration of PX DNA dissolved in 1X TAE.

**RNA isolation and quantitative RT-PCR (qRT-PCR)**

Total RNA was extracted using RNEasy mini kit (Qiagen) by following the manufacturer's instructions.

cDNA synthesis was carried out using the iScript cDNA Kits (Bio-Rad) as instructed. Then, qRT-PCR was carried out using Sybr green PCR master mix (Bio-Rad) in a Bio-Rad thermal cycler using Myogenin (Myog) and Myosin Heavy Chain  MHC) specific primers. GAPDH primer pairs were used as housekeeping gene for normalizing the values of Myog and MHC.

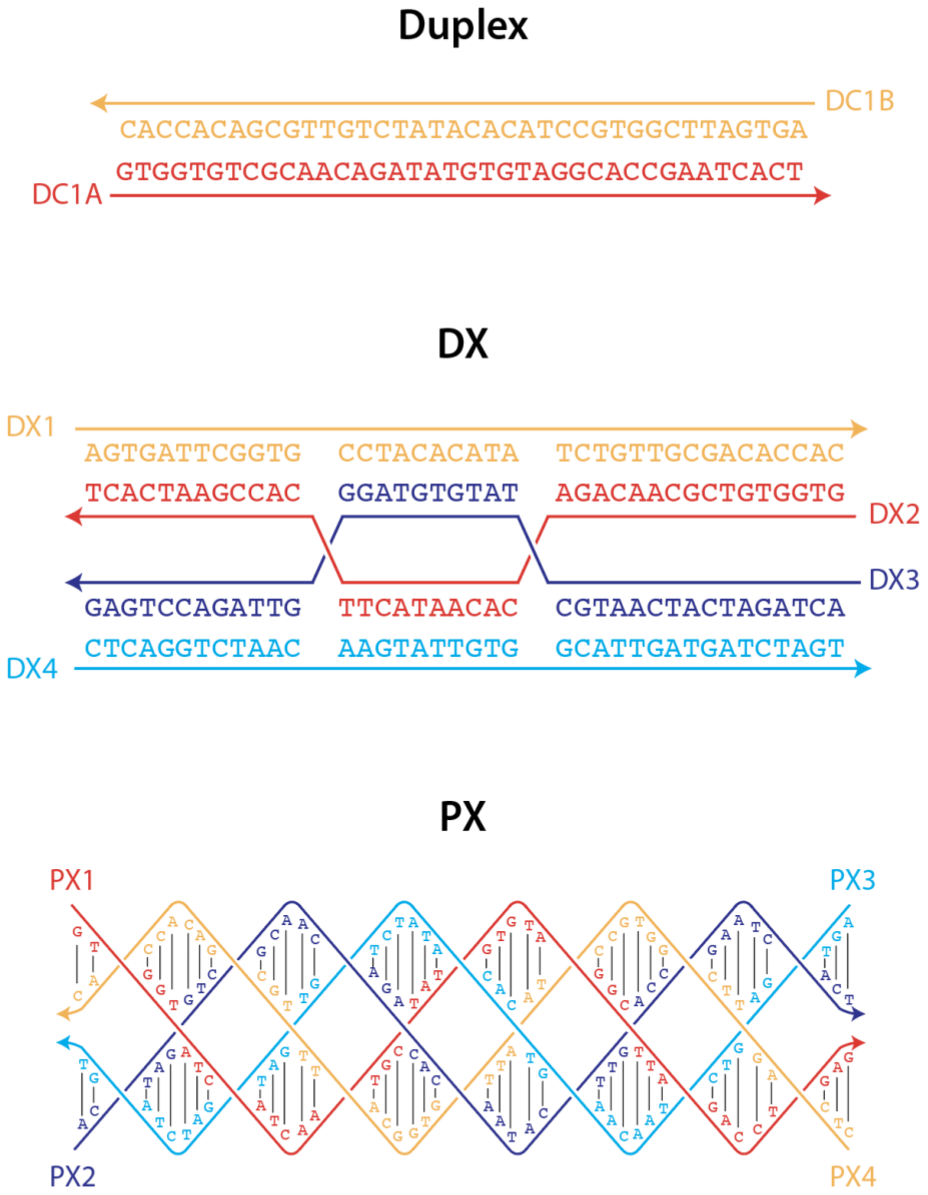


**Figure S1**. Design and sequences of duplex, DX and PX DNA motifs used in this study. Arrows indicate 3' ends of strands.

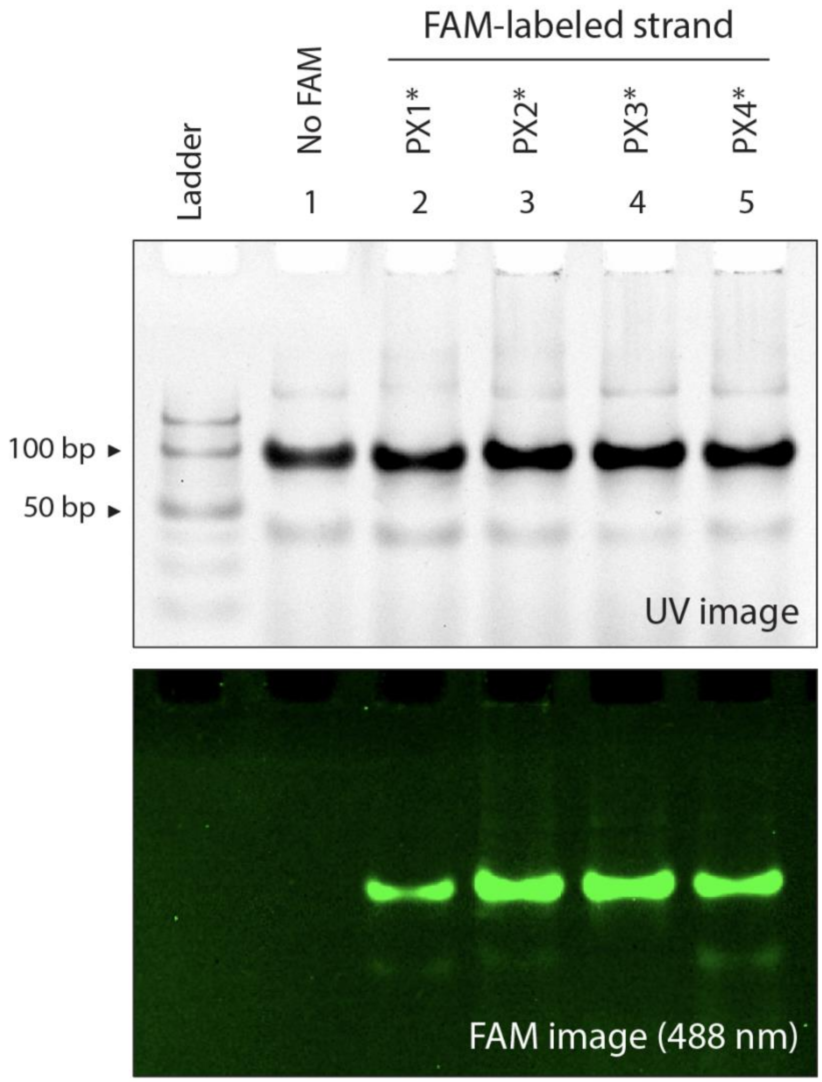


**Figure S2**. Analysis of PX formation using fluorophore-tagged component strands. PX complexes were annealed with only one of the four component strands labeled with fluorescein (FAM). Non-denaturing PAGE showed that all four strands are present in the complex (lanes 2-5) compared to a non-labeled complex (lane 1).

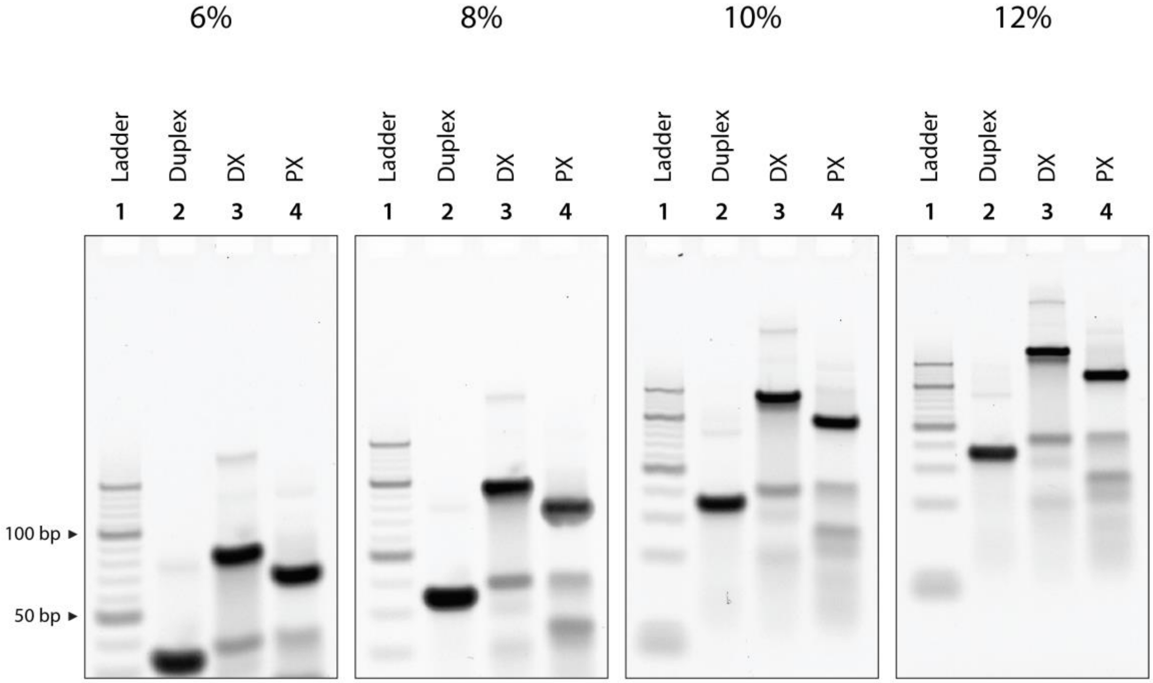


**Figure S3**. PAGE gels for Ferguson plot analysis of the three structures (duplex, DX and PX). Slopes from the Ferguson plot show that the PX motif is similar to a DX and distinct from double stranded B-DNA.

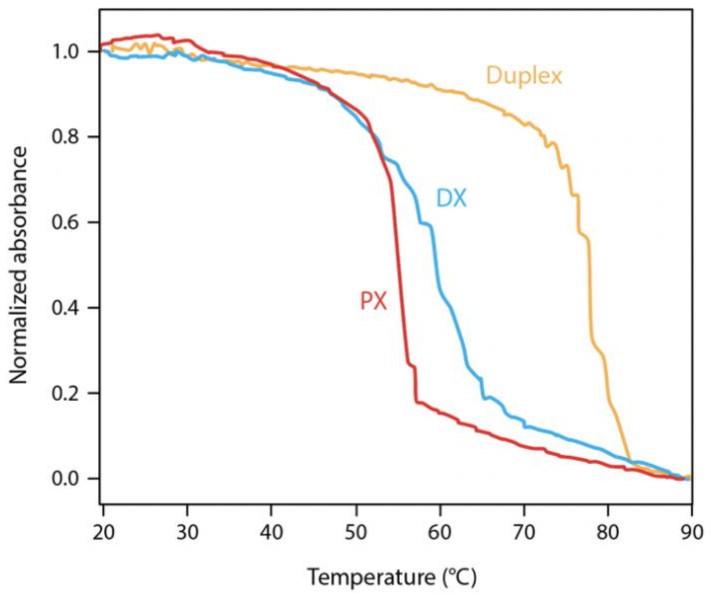


**Figure S4**. UV melting profiles of the duplex, DX and PX structures used in this study.

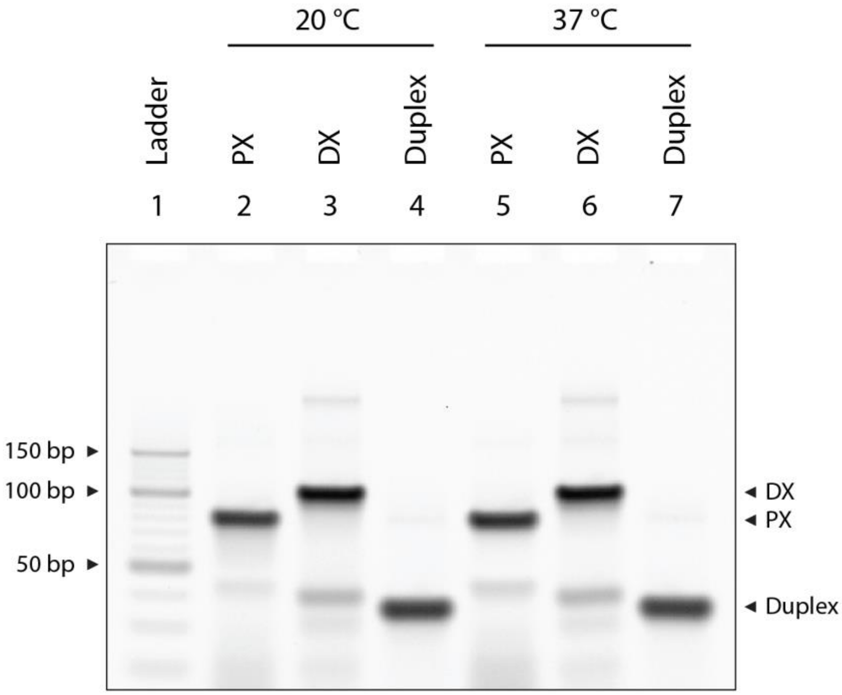


**Figure S5**. Duplex, DX and PX motifs are intact at 37 °C for at least 24 hours.

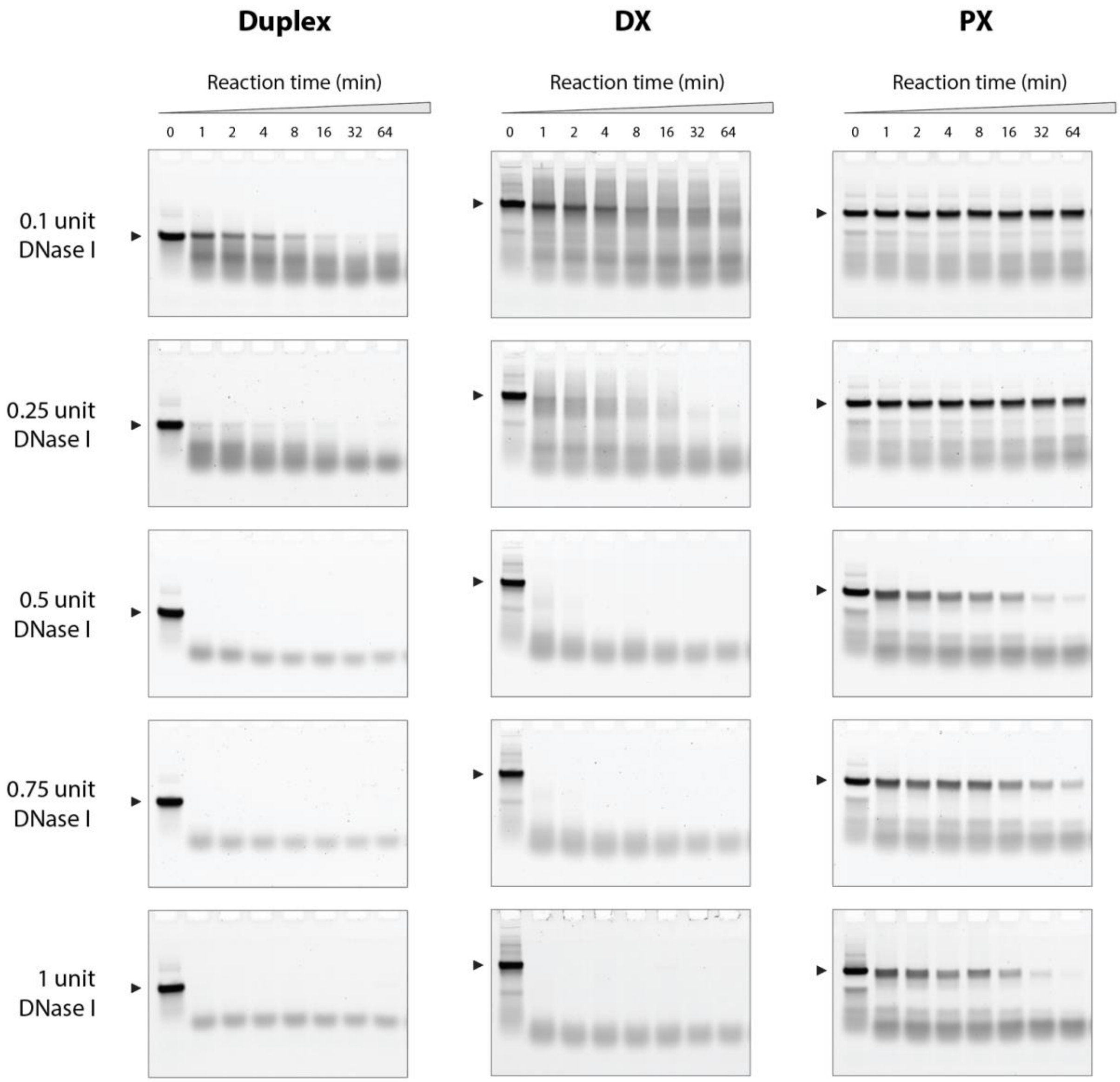


**Figure S6**. DNase I treatment of duplex, DX and PX motifs. Non-denaturing PAGE showing degradation of the three structures when incubated with different amounts of DNase I enzyme.

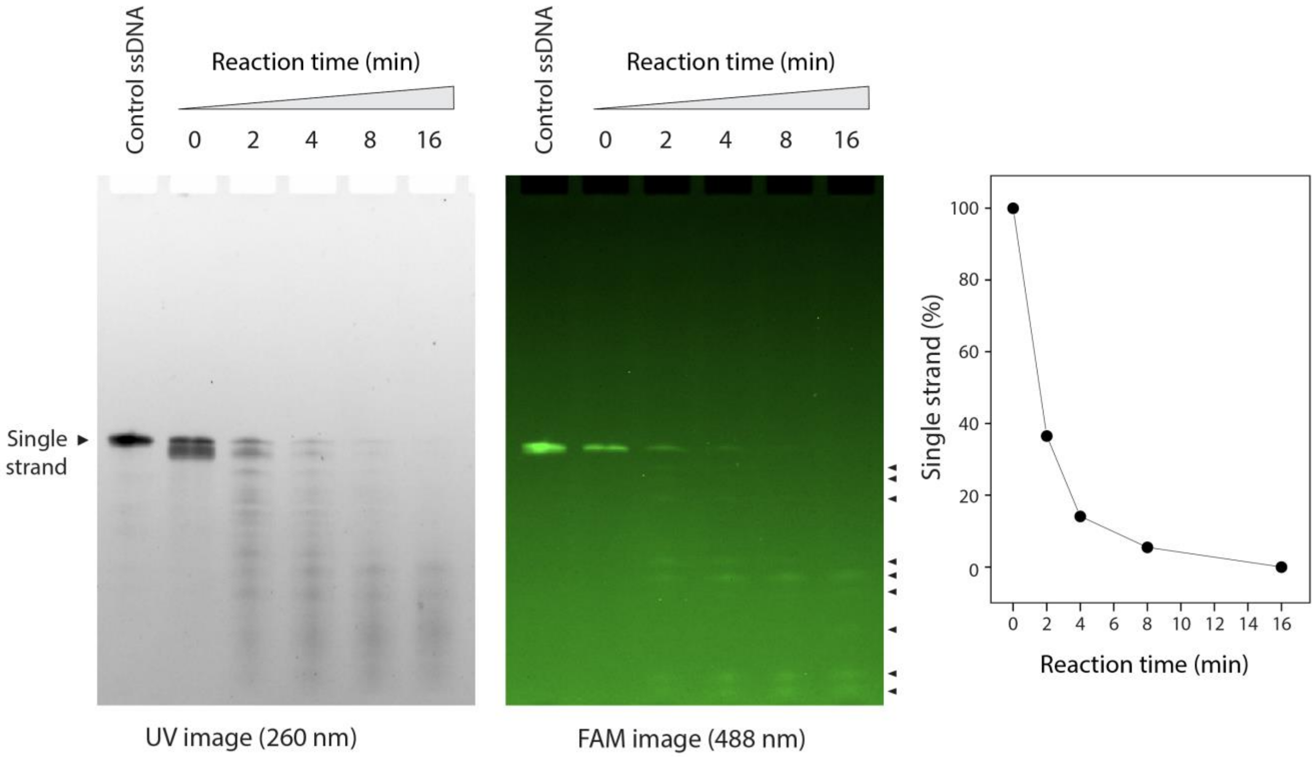


**Figure S7**. Non-denaturing gel showing DNase I reaction products from PX DNA at different reaction times. Strand 2 in the PX was FAM-labeled and imaged at 488 nm (right).

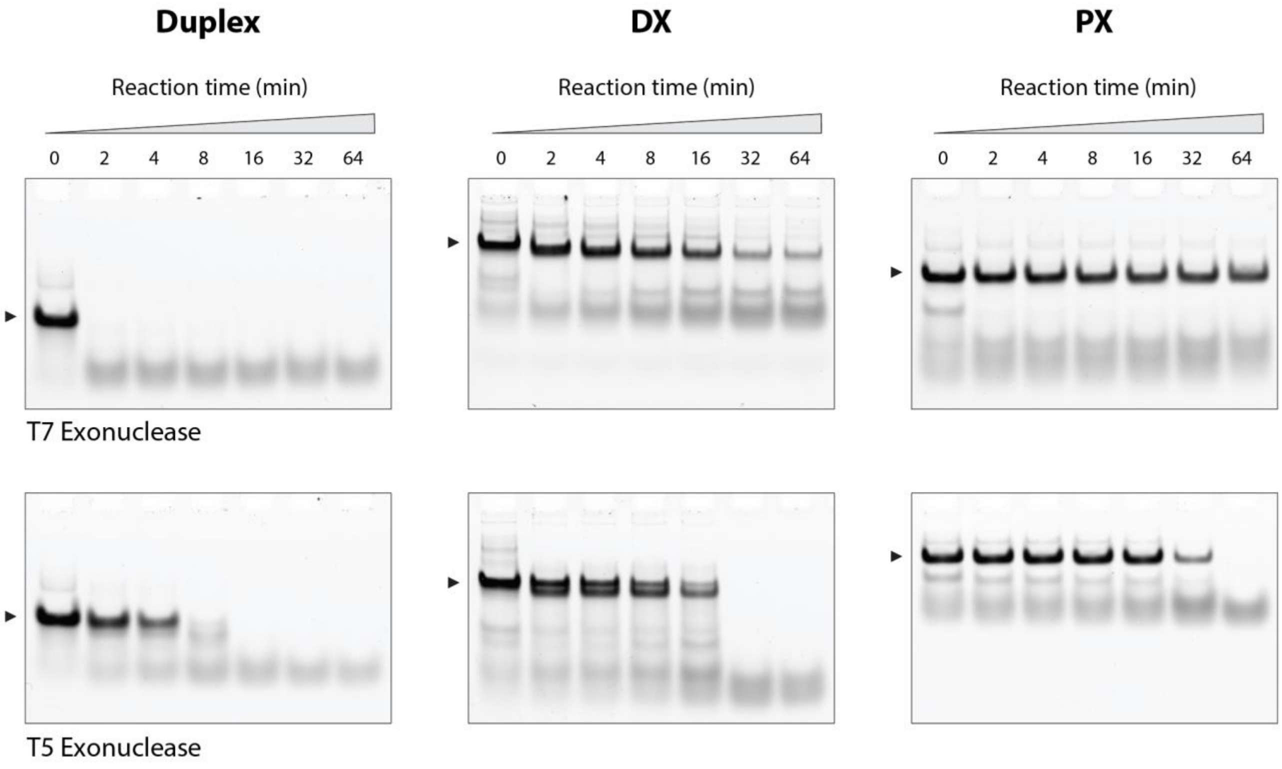


**Figure S8**. Nuclease resistance of tested structures against T7 and T5 exonucleases (full gels of images in Figure 2e-2f).

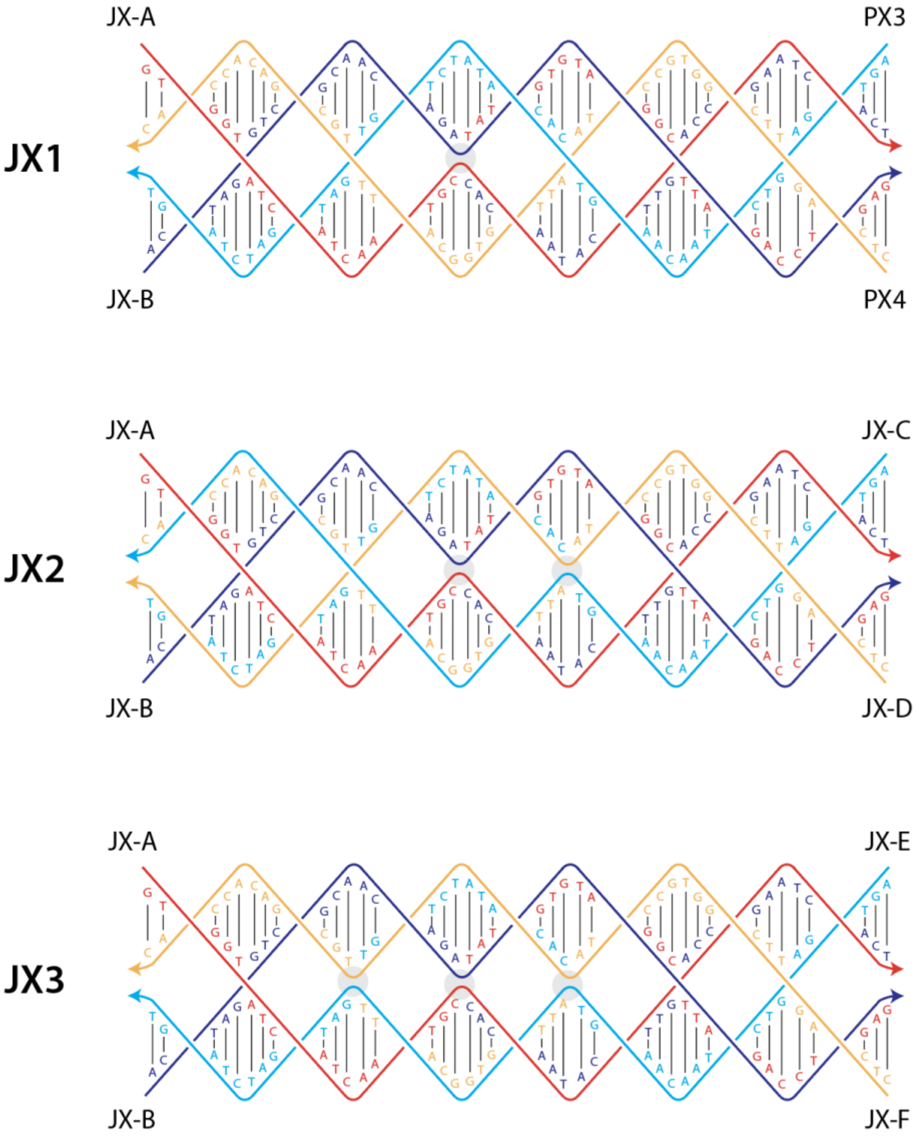


**Figure S9**. Design and sequences of the JX$_n$ DNA motifs used in this study. Arrows indicate 3’ ends of strands.

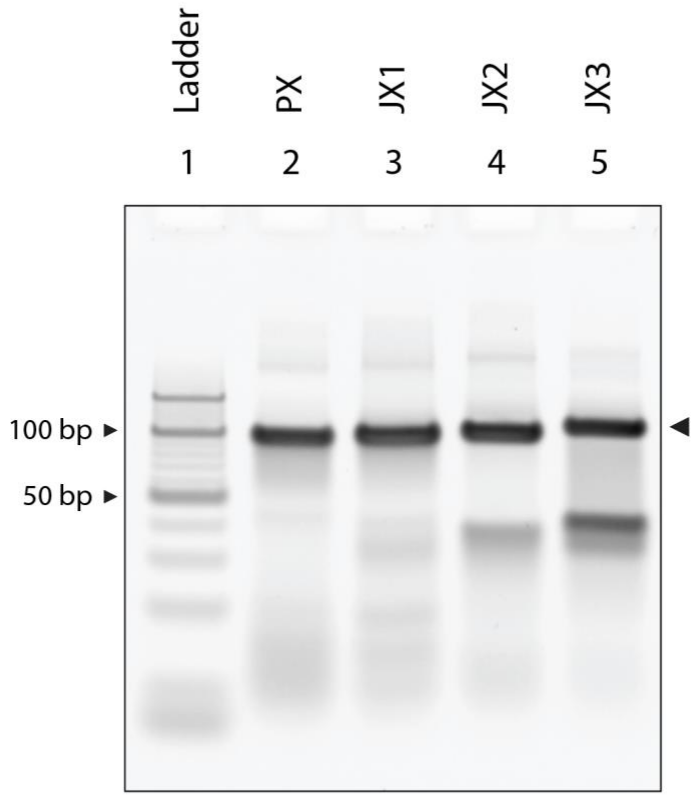


**Figure S10**. Non-denaturing PAGE showing the formation of $JX_n$ structures.

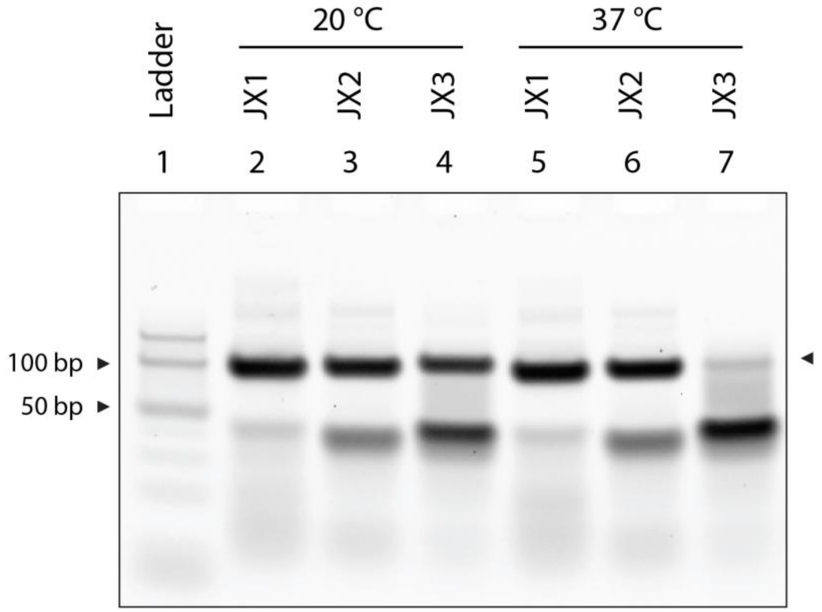


**Figure S11**. Stability of $JX_n$ motifs at 37 °C for 2 hours. $JX_1$ and $JX_2$ structures are stable at 37 °C while the $JX_3$ motif is degraded.

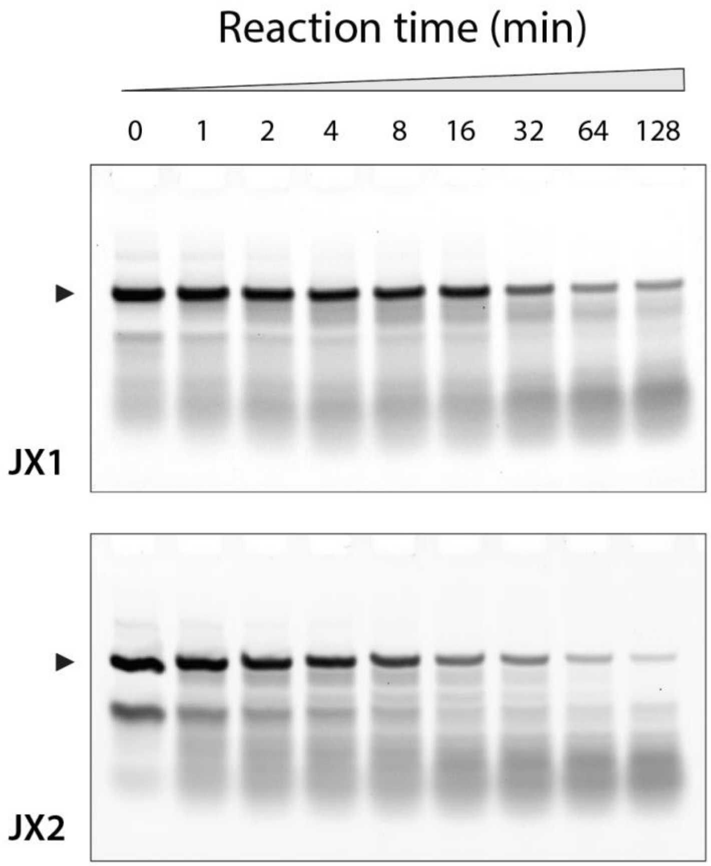


**Figure S12**. Non-denaturing PAGE showing degradation of $JX_1$ and $JX_2$ structures when incubated with 0.1 unit DNase I enzyme.

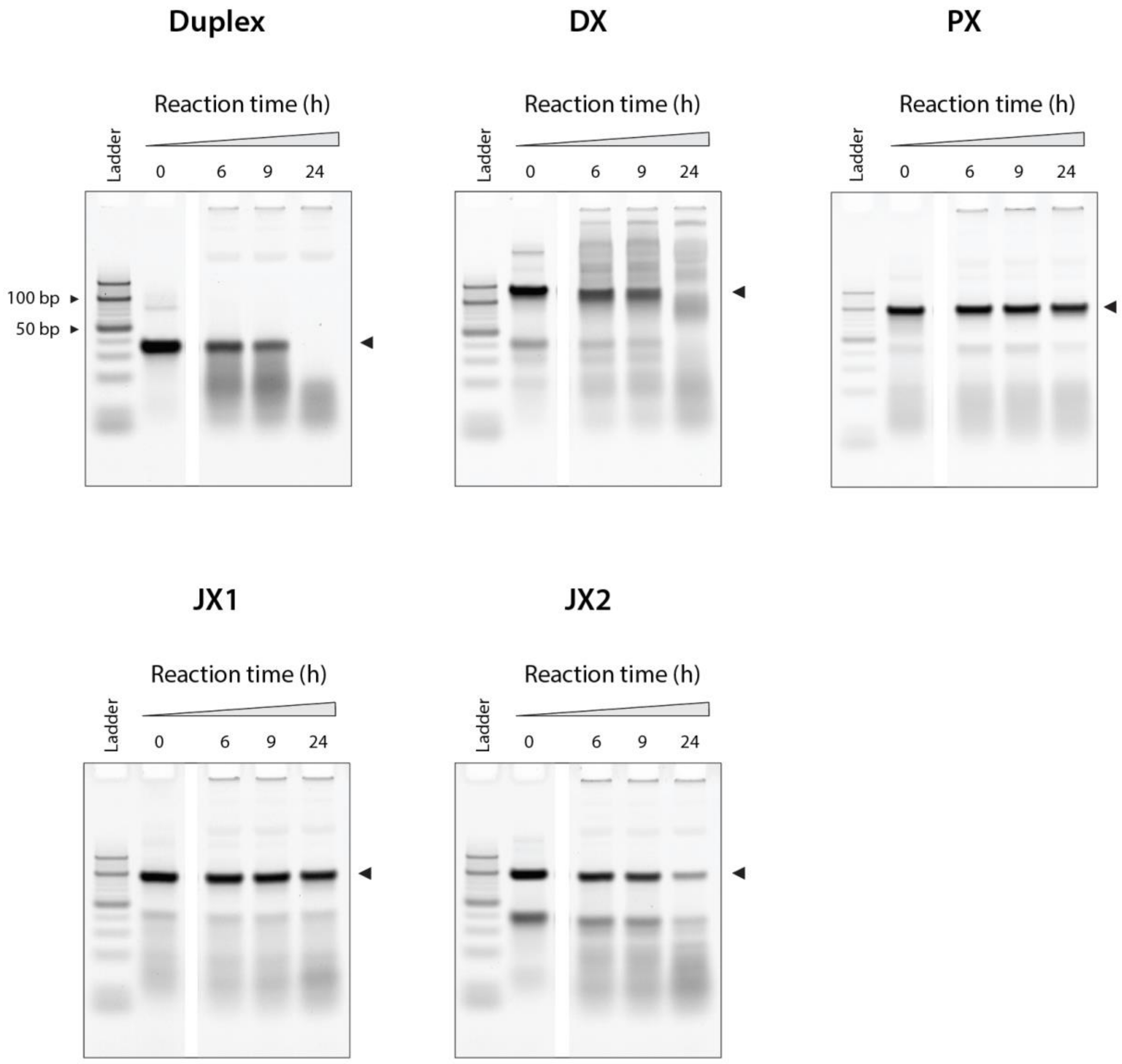


**Figure S13**. Full gels showing structures incubated in 10% FBS (also shown in Figure 4 in main text).

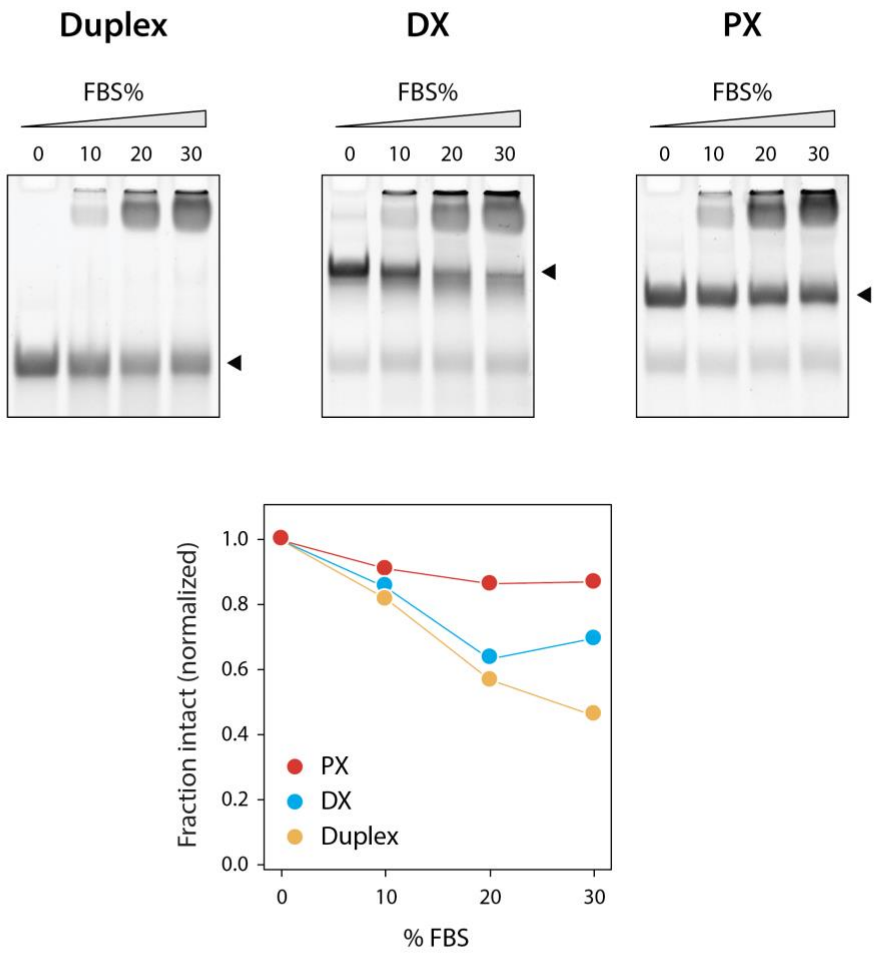


**Figure S14**. Stability of duplex, DX and PX motifs incubated in different amounts of FBS.

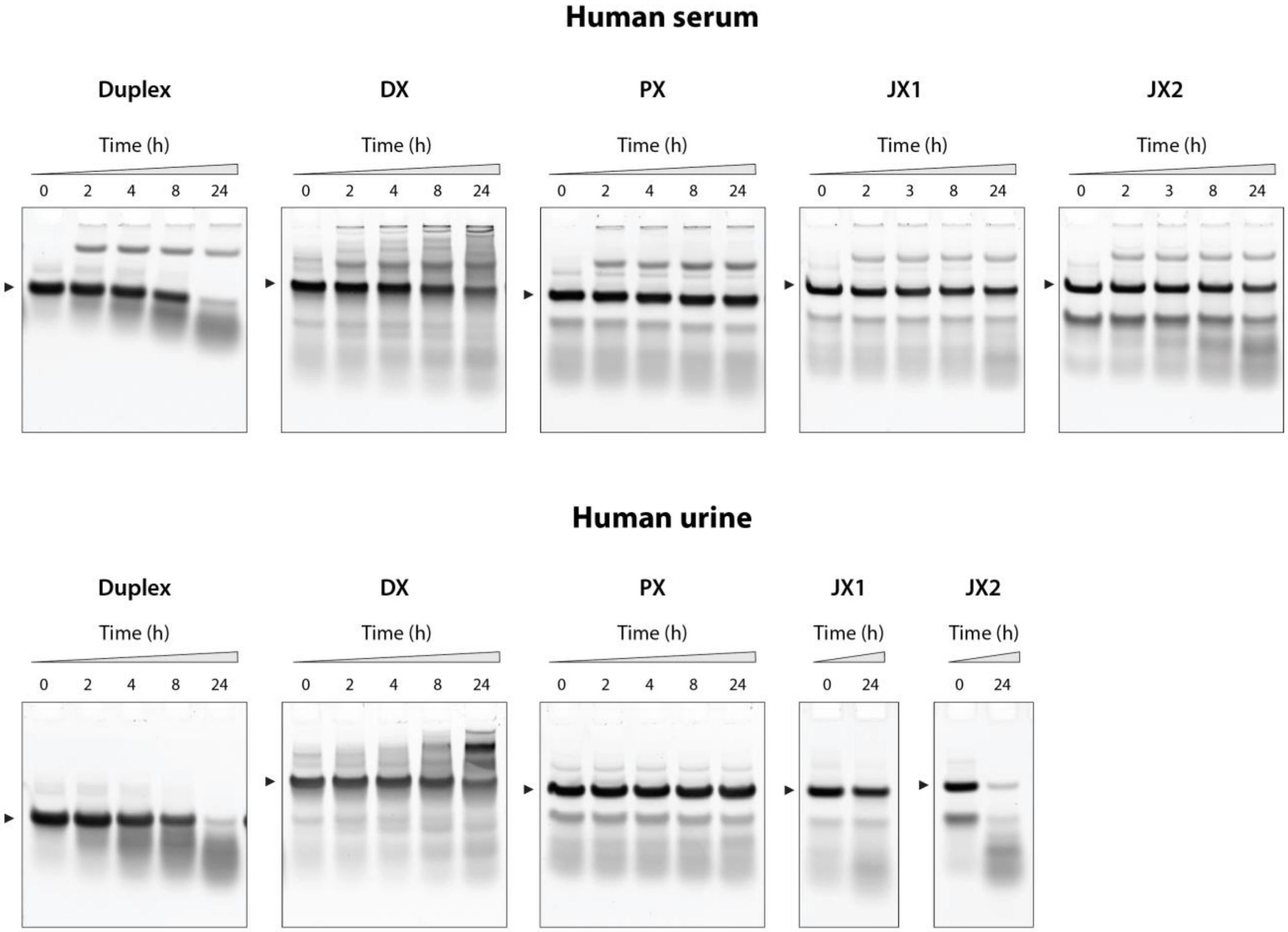


**Figure S15**. Stability of structures incubated in different amounts of human serum (top) and human urine (bottom).

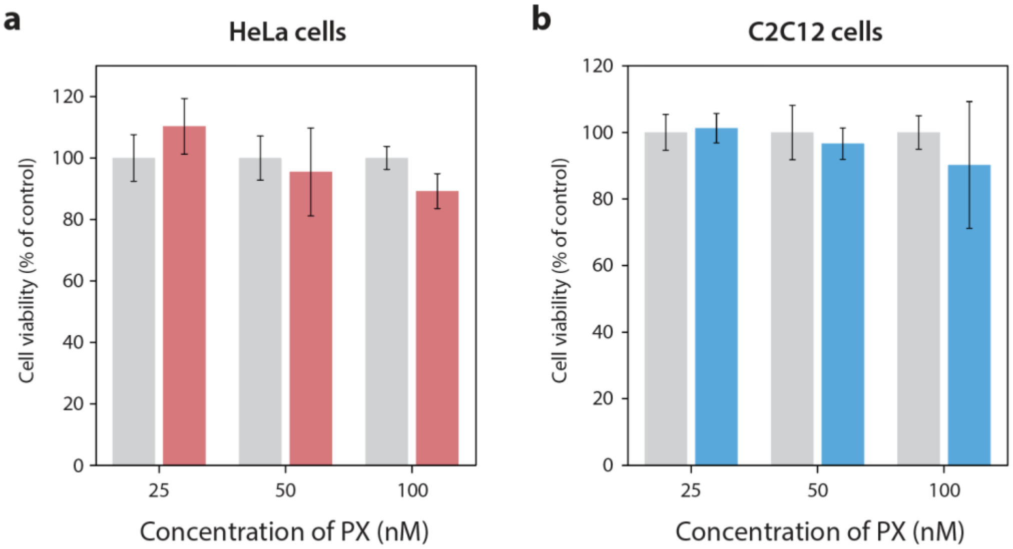


**Figure S16**. MTT assay showing cell viability after 48 hours in the presence of different concentrations of PX in (a) HeLa and (b) C2C12 cells.
